# The speed and phase of locomotion dictate saccade probability and simultaneous low-frequency power spectra

**DOI:** 10.1101/2023.06.22.546202

**Authors:** Lydia Barnes, Matthew J Davidson, David Alais

## Abstract

Everyday we make thousands of saccades and take thousands of steps as we explore our environment. Despite their common co-occurrence in a typical active state, we know little about the coordination between eye-movements and walking behaviour and related changes in cortical activity. Technical limitations have been a major impediment which we overcome here by leveraging the advantages of an immersive wireless virtual reality (VR) environment with three-dimensional position tracking, together with simultaneous recording of eye-movements and mobile electroencephalography (EEG). Using this approach with participants engaged in unencumbered walking along a clear, level path, we find that the likelihood of eye-movements at both slow and natural walking speeds entrains to the rhythm of footfall, peaking shortly after the heel-strike of each step. Simultaneous EEG recordings reveal a concomitant modulation entrained to heel-strike, with increases and decreases in oscillatory power for a broad range of frequencies. The peak of these effects occurred in the theta and alpha range for both walking speeds. Together, our data show that the step-rate of locomotion influences other behaviours such as eye movements and produces related modulations of simultaneous EEG following the same rhythmic pattern. These results reveal gait as an important factor to be considered when interpreting saccadic and time-frequency EEG data in active observers.

## Introduction

Perception in real-world contexts often takes place in the service of actions executed to achieve behavioural goals. Actions such as visual search, reaching and locomotion all require perception and attention so that they can be effectively executed. Our most frequent action is to move our eyes, and we make upwards of 150,000 saccades per day (Carpenter, 1988, 2000), but we also take several thousand steps each day as we move about our environments. We are therefore inherently active beings, which necessitates the study of perception and related behavioural and neural effects during activity. Despite this, most of what we know comes from studies in which participants are immobile and required to suppress head and eye movements. A number of recent studies, however, have begun to study vision and related behaviours in more active contexts (Cao et al., 2020; Cao & Händel, 2019; Chen et al., 2022; Gramann et al., 2014, 2021; Gwin et al., 2011).

One impetus behind the push to study vision in active contexts comes from relatively recent work showing that locomotion boosts cortical responsivity in the rodent primary visual cortex (Kaneko et al., 2017; Vinck et al., 2015). These effects of locomotion are robust and widely reported (Erisken et al., 2014; Fu et al., 2014; Lee et al., 2014; Mineault et al., 2016; Niell & Stryker, 2010), for a review, see: (Parker et al., 2020)) and studies have begun to appear in recent years exploring the role of activity on perception and behaviour in human subjects. In most of these studies, activity conditions are produced either by peddling on a stationary exercise bicycle or by walking on a treadmill. Despite the clear-cut results in rodent studies, findings from these studies have produced some mixed results. In one walking study, (Benjamin et al., 2018) measured both electroencephalography (EEG) and visual contrast psychophysics and found no evidence for an increase in either spontaneous firing rate or contrast sensitivity when walking on a treadmill. (Cao & Händel, 2019) on the other hand, measured EEG and visual performance in free-walking participants and found that walking boosted peripheral visual processing. The evidence supporting this was a relative reduction in the SSVEP amplitude evoked from a central target during walking compared to standing, and behavioural data showing greater surround-suppression on central target detection during walking.

Other studies have investigated the influence of physical activity on visual responses by measuring EEG while participants pedalled on stationary exercise bicycles. These studies have generally reported a sensory and perceptual enhancement during activity. (Bullock et al., 2015) found faster reaction times in an odd-ball detection task during exercise, although found no change in detection performance. EEG results showed the P1 component evoked by odd-ball targets was larger and peaked earlier in low-intensity exercise (relative to stationary and high-intensity conditions). (Bullock et al., 2017) inferred feature-selectivity from scalp recordings using an inverted encoding model. They found narrower bandwidth for low-intensity cycling relative to high intensity or stationary conditions, and an equivalent improvement in orientation discrimination across both exercise conditions relative to stationary. In a similar study, (Dodwell et al., 2021) found faster reaction times in a visual search task during low- and high-intensity exercise and evidence from ERPs of reduced distractor interference.

In the present study, we combine several features from the studies reviewed above, while incorporating new elements. In preference to a treadmill or exercise bicycle, we had participants walk freely along a short path in wireless virtual reality (VR) while viewing a stream of visual stimuli and performing an oddball detection task. While walking and performing this task, participants also wore a mobile EEG system designed to fit with the head-mounted display, enabling the continuous acquisition of eye-movements and neural activity. By resampling behaviour and neural activity into sequential steps and strides, we were able to show that specific phases of the step-cycle alter continuous gaze direction and spectral dynamics of EEG. To preview the results, we find clear modulations within the gait cycle of the likelihood of saccade onset - despite targets being presented at a constant rate during locomotion. During both slow and natural walking speeds, periods of increased and decreased power are evident in the time-frequency transform of EEG data, with peak frequencies for these effects in the theta and alpha bands.

## Methods and materials

### Participants

We recruited 37 healthy volunteers (14 male, age 21.81 ± 5.29). Participants were fluent English speakers and had normal or corrected to normal vision. The final sample consisted of 19 participants (6 male, age 21.05 ± 3.43). This study was approved by the University of Sydney Human Research Ethics Committee (HREC 2021/048), and all participants received either course credit or 20 AUD per hour of their time.

### Apparatus and virtual environment

The virtual environment was built in Unity (version 2020.3.14f1) incorporating the SteamVR Plugin (ver 2.7.3; SDK 1.14.15), on a DELL XPS 8950, with a 12th Gen Intel Core i7-12700K 3.60 GHz processor, running Microsoft Windows 11. The virtual environment consisted of a light green sky and green ground plane. Two overlaid walkways stretched across the ground plane. A longer walkway, in light red, extended seven metres. Superimposed on the centre of this walkway sat a shorter walkway, in dark red, extending half that distance.

Participants walked along this simulated seven metre (or three and a half metre) track wearing the HTC Vive Pro Eye with integrated head-mounted display (HMD) and a wireless adapter kit (130 gram weight). Two wireless hand-held controllers were carried to collect participant responses using a right index finger trigger button. The HMD houses two 1440 x 1600 pixel (3.5” diagonal) AMOLED screens, with a 110 degree field of view refreshed at 90 Hz. We recorded gaze origin and gaze direction information using the integrated Tobii eye-tracking technology and SRanipal SDK (v1.1.0.1). Five HTC Base Stations (v 2.0) were used to record the three-dimensional coordinates of the HMD, gaze-origin and gaze-direction at 90 Hz resolution. Participants also wore a HMD-compatible mobile EEG system (DSI-24, Wearable Sensing), which wirelessly transmitted data to Wearable Sensing’s bespoke acquisition software running on a Microsoft Surface tablet. Triggers to mark task events in the EEG data were transmitted from the stimulus computer to a NeuroSpec MMBT-S trigger box, which took serial input via a USB cable and output triggers via a D-SUB 25 parallel port cable. The signal was routed through Wearable Sensing’s trigger box before being wirelessly transmitted to the EEG headset. EEG Data and trigger information were saved to the tablet for offline processing.

### Procedure

Participants were first invited to read the information and ask questions before giving informed consent. Next, participants were asked to sit while a tester fit a mobile EEG system to their head. The system (DSI-24, Wearable Sensing) consists of a rigid frame with adjustable straps around and across the head. The tester centred the system on the participant’s head and tightened it for stability. Participants were asked to identify any pressure points, and the tester adjusted tightness accordingly. The tester then rotated the dry electrodes so that they pushed through the participant’s hair. The electrodes consisted of a ring of small teeth on a rigid arm, with a small spring between the electrode and the rigid arm for flexible fit. The tester adjusted the electrodes until the teeth sat cleanly against the scalp, with an impedance below 10 kΩ. Two supplementary flat electrodes were attached to the earlobes with clips. Participants were invited to view their EEG and observe the effect of face movements on the data.

Once the EEG system was in place, the tester fit the HMD over the top. The HMD attached to the EEG system with velcro straps, which were tightened for stability and comfort. Some participants’ head shape required that the frontal strip of EEG electrodes be moved up to allow the HMD to fit snugly against their face. The tester then strapped the HMD’s wireless adapter to the top of the participant’s head, and gave them a hand-held controller.

Participants were shown the virtual environment and given time to familiarise themselves with it, while the tester ran an eye tracking calibration. Participants were informed that they could safely move along the red walkways in the virtual space, and that the experimenter would always be present to ensure they were far from obstacles in the testing room.

Participants began with five static practice trials. They were told what would happen in a trial, and given time to ask questions. Each practice trial began with a warning, the text “Ready?”, presented centrally in front of the participant for 300 ms. This was followed by a fixation cross presented alone for 500 ms, and a five second rapid serial visual presentation (RSVP) stream. The stream consisted of 25 images, 24 of them unique within the trial, one of them (the target) a repetition of the image before. Each image was displayed for 200 ms. Images could appear at any one of eight locations arranged in two quadrants, close to or far from fixation. The fixation cross remained in place throughout the stream. Participants were instructed to pull the trigger button under their index finger as quickly as possible when they saw a repetition, that is, the target image. At the end of the stream, they had three and a half seconds to rest before the next “Ready?” warning. The tester guided the participant through the practice trials and confirmed that they understood the basic trial structure before moving on. The tester then explained the main task, and started the main testing session.

The main session consisted of three blocks of 100 trials each. Each block corresponded to one condition: static, slow walking, or natural walking. The static condition was identical to the practice trials described above. In the two walking conditions, the stimulus content and timing were the same as in the static condition. Here, however, participants were asked to walk a prescribed distance during the RSVP stream. In the slow walking condition, participants walked the length of the three and a half metre dark red walkway. In the natural walking condition, participants walked the length of the seven metre light red walkway. This was designed to suit a 120 bpm walking speed with a 65 cm step length over the five second RSVP stream (6.5 m + 50 cm buffer). Participants were invited to sit, rest, and loosen the HMD in between blocks to minimise eye strain.

### Stimuli

Stimuli were taken from the THINGS image database (Hebart et al., 2019). We selected thirty concepts with high concreteness and frequency ratings: arm, baby, ball, bean, bed, bird, camera1, cup, dog, ear, finger, fish, flower, gun, hair, horse, leaf, lion, missile, paper, pig, rifle, sand, snake, spider, syringe, tiger, toilet, tree, worm. For each concept, we selected 10 exemplars creating a stimulus set of 300 images. Each image was presented eight times within a condition (24 unique images per trial x 100 trials), excluding presentations as the target image.

Stimuli were presented at a distance of 1 m in front of the participant in the virtual space, in one of eight locations. Four of these locations were close to the centrally presented fixations, with the centre of the image four degrees of visual angle away from the centre of the fixation cross, diagonally up-left, up-right, down-right, or down-left. The remaining four locations were similarly placed on the diagonal, but peripherally, with the centre of the image 12 degrees of visual angle from the centre of the fixation cross. Stimuli were 12.25 cm tall and wide, subtending seven degrees of visual angle.

### Eye movement data

Eye movements were calibrated at the start of each experiment using the manufacturers 4-point calibration procedure. Additional calibration was performed after any change to the headset (such as after breaks, between blocks). Continuous recordings of gaze origin and gaze direction were coordinated in 3D coordinates (each frame of all trials at 90 Hz resolution), the following preprocessing steps were used to identify microsaccades following the algorithm of (Engbert & Kliegl, 2003).

First, whole-trial time-series of the eye-position and eye-origin data were tested for outliers. Blinks were identified on the basis of outliers in the first-derivative of these time-series that exceeded ± 0.8 m. Raw time-series data was then linearly interpolated from −100 ms before to +100 ms after each blink. Identical windows were interpolated for the gaze origin and gaze direction time-series on the x, y, and z axes. After blink interpolation, we applied a two-dimensional velocity-based algorithm to identify saccades within each trial. Time series of eye-position data were transformed to velocities and smoothed with a 5 sample (55 ms) moving average to suppress noise. Microsaccades were then detected as outliers exceeding velocity thresholds that were computed separately for the x and y time series. Following Engbert & Kliegel (2003), we multiplied the median estimator by 6 x the standard deviation of each velocity time series, to protect each threshold computation from noise. As each threshold in this algorithm is computed relative to the standard deviation on each trial, it is robust to different noise levels between trials, conditions and participants (Engbert & Kliegel, 2003). We retained the onset, duration and raw position of each microsaccade that was detected for allocation into our stride-based epochs, detailed below.

### Gait extraction from head-position data

We applied a peak-detection algorithm to the time-series of vertical head position data to estimate the phases of the stride-cycle. As the vertical centre of mass follows near sinusoidal changes during walking, troughs correspond to when both feet are placed on the ground during the double support stance phase (Gard et al., 2004; Hirasaki et al., 1999; Moore et al., 2001; Pozzo et al., 1990). We epoched all individual steps based on these troughs, and normalised step-lengths for stride-cycle analysis by resampling the time-series data to 200 data points (0-100% stride-cycle completion, increment of 0.5%). After peak-detection, each trial was also visually inspected to identify trials for exclusion, based on wireless signal drop-out or poor gait extraction.

### Cycles-per-stride analyses

We assigned target- and saccade-onsets to their position in the simultaneously occurring stride cycle (i.e. a percentile from 1-100 % stride-completion).We then averaged target onsets and saccade counts within 40 linearly spaced bins (with zero overlap) and applied Fourier fits to detect the presence of oscillations.

Significant oscillations were detected via a two-step procedure. We first fit a sequence of Fourier series models within a forced frequency range to the observed group-level data. These models stepped from 0.2 to 10 cycles per stride, in 0.2 increments, using MATLAB’s curve fitting toolbox and the equation:

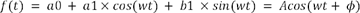

Where *a0* is a constant, *a1* and *b1* are cosine and sine coefficients, *t* is time and *w* is the periodicity per stride-cycle. *A* and ¢ are the resulting amplitude and phase for the sinusoidal fit. For each forced fit at each frequency, this routine implemented a non-linear least squares method (400 iterations), from which we retained the goodness of fit (*R*^2^) as our critical value. As a second stage, we assessed the likelihood of these fits occuring by chance by comparing them to a null distribution obtained from a non-parametric shuffling procedure. For each participant the data at a given percentile was selected from another percentile at random (without replacement), and the group-level fitting procedure was repeated as described above. The *R*^2^ values from 1000 permutations of this procedure were used to quantify the upper 95% Confidence Interval (CI) of the null distribution at each frequency, and the presence of a significant oscillation was inferred when the observed *R*^2^ value exceeded this upper bound.

### EEG recording and pre-processing

Electroencephalography was recorded with a dry headset (DSI-24, Wearable Sensing) from 19 standard electrode positions corresponding to the international 10-20 system (Fp1, Fp2, F7, F3, Fz, F4, F8, T7, C3, Cz, C4, T8, P7, P3, Pz, P4, P8, O1, O2), and two linked earlobe electrodes (A1, A2) for offline re-referencing. Electrode impedances were kept below 20 kΩ during recording, sampled at 300 Hz and referenced online to Pz. During pre-processing, EEG were re-referenced to the average of A1 and A2, and bandpass filtered between 0.1 and 40 Hz prior to epoching per trial. Bad channels were removed with pyprep deviation and correlation metrics. Each trial was epoched −100 ms to 6500 ms relative to trial onset, and baseline corrected using the 100 ms pre-trial period. Across participants, a total of 11 trials were dropped during epoching (maximum two per participant) due to a data drop-out in the epoch.

### Gait-cycle based EEG

For gait-cycle based EEG analyses, each step was epoched from −600 ms to +600 ms relative to step cycle onset and completion, respectively. Outliers were identified for exclusion when the epoch standard-deviation exceeded 1.5 interquartile ranges above or below the upper and lower quartiles of the distribution of epoch standard deviations across all trials. This method removed an average of 11.5 (*SD* = 17.5) and 37.2 epochs (*SD* = 34.8) in the slow and natural walking speeds across subjects. To investigate time-frequency activity relative to position in the gait-cycle, we convolved our time-series signal with a set of complex Morlet wavelets. Each Morlet wavelet was defined by a complex sine wave tapered by a Gaussian, and ranged from 3 Hz to 40 Hz in 50 linearly spaced steps. As wavelet peak frequency increased, the full-width at half-maximum ranged from 600 to 100 ms (Cohen, 2019). After conversion to the time-frequency domain, single-trial gait-epochs were resampled from 1-100% cycle completion to allow comparisons, with an additional +-30% buffer (260 data points, in increments of 0.5%). Due to the cyclic nature of locomotion, time-frequency power within each frequency was normalised to percent change over the entire epoch using the full epoch as a baseline.

### Saccade-locked EEG

We epoched saccade-locked EEG activity from −600 to +1100 ms relative to saccade onset. Saccade epochs which began before RSVP onset, or exceeded the trial duration were omitted from analysis. We applied the same outlier-detection and Morlet-wavelet convolution as described above, with no additional resampling in units of time. To facilitate comparisons, saccades were additionally sorted based on their occurrence within a co-occurring step cycle into four quartiles (1-25%, 25-50%, 51-75%, 76-100% step cycle completion). All saccade locked time-frequency power was normalised to percent change over the entire epoch, using a baseline computed from the average of all saccades (irrespective of gait quartile).

For all EEG analyses we corrected for multiple comparisons using non-parametric cluster-based permutation tests (Maris & Oostenveld, 2007). For time-frequency data, contiguous *t*-scores from dependent samples *t*-tests were retained which exceeded *p* < .01 (uncorrected), and compared to null-distributions obtained using Monte Carlo permutation tests (2000 repetitions). Each permutation exchanged data labels at random, and the maximum sum of contiguous *t*-scores was retained to create the null-distribution of clustered test-statistics (Maris & Oostenveld, 2007). By comparing the observed cluster statistic to the empirical cumulative of the null distribution we finally obtained the cluster-level p-value (reported here as *p*_cluster_) corrected for multiple-comparisons.

### Exclusions

Thirty-seven participants took part in the study. One participant found the set-up uncomfortable, and finished before the task was completed. Two participants’ behavioural data were not properly recorded. Three were excluded for substantial EEG drop-out, seven for EEG trigger failure. Finally, five were excluded for missing eye tracking data, leaving 19 participants with full EEG, eye-tracking, and behavioural data. All results reported below are from these 19 intact datasets.

## Results

We designed a RSVP task for participants to view while walking at slow and natural walking speeds in a wireless VR environment. Below, we focus on the characteristics of the EEG and eye-movement data simultaneously recorded during this task (Figure 1). The results relevant to object categorisation will be reported in another manuscript in preparation.

**Figure 1.**
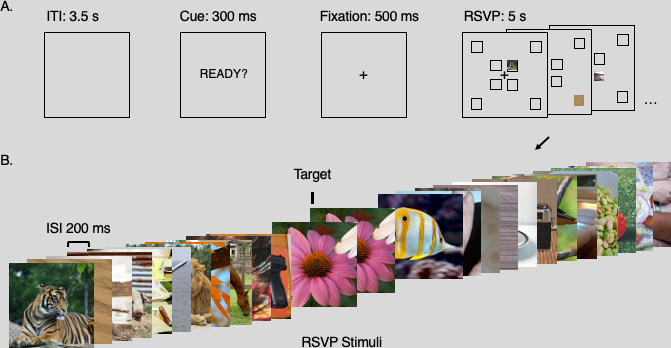
Paradigm. Panel A shows the core components of the trial. Trials were separated by a substantial gap to allow participants time to adjust their position between walking lengths of the room. A cue warned participants to settle into their start position. This was followed by a brief fixation cross before the rapid stimulus stream. Stimuli could appear in any one of eight positions, as marked on the figure (the square location markers for illustration only). Panel B shows an example set of stimuli for a trial. Each trial included 25 images, 24 of them unique, one of them a target, presented at 5 Hz. Each image was presented for 100 ms, with a blank gap of 100 ms between images. Participants were instructed to press a button whenever they saw an image presented twice in succession (the second presentation is referred to as the target).

### Walking speed alters gait and gaze dynamics

As expected, the instructions to walk at a natural or slow walking speed resulted in differences in gait parameters and gaze direction. When walking slowly, participants on average lengthened the duration of their steps and exhibited a decrease in the sinusoidal amplitude of changes to the centre of mass over time (Figure 2). Figure 2A displays an example trial from one participant at each walking speed, exemplifying the elongated step duration during slow walking conditions. Figure 2B displays the grand average histograms of step durations for all participants in each condition (*N*=19). On average, our peak detection algorithm recorded a total of In order to compare stride-cycle dynamics across walking speeds, we applied a stride-cycle resampling procedure common to gait and posture research. This method resamples the duration of all strides (two-sequential steps) between 1–100% stride-cycle completion, a normalisation enabling comparison of performance measures relative to the phase of locomotion. Figure 2C displays the grand-average detrended head height per stride cycle at slow and natural walking speeds for our participant*s*. Approximate locations of swing and stance phases, as well as heel-strike and toe-off are indicated. These are based on prior studies which have measured foot pressure and head acceleration simultaneously (Mulavara & Bloomberg, 2002) and define the heel-strike and toe-off points as 10% of step-cycle period before and after mid-stance, respectively (MacNeilage & Glasauer, 2017).

**Figure 2.**
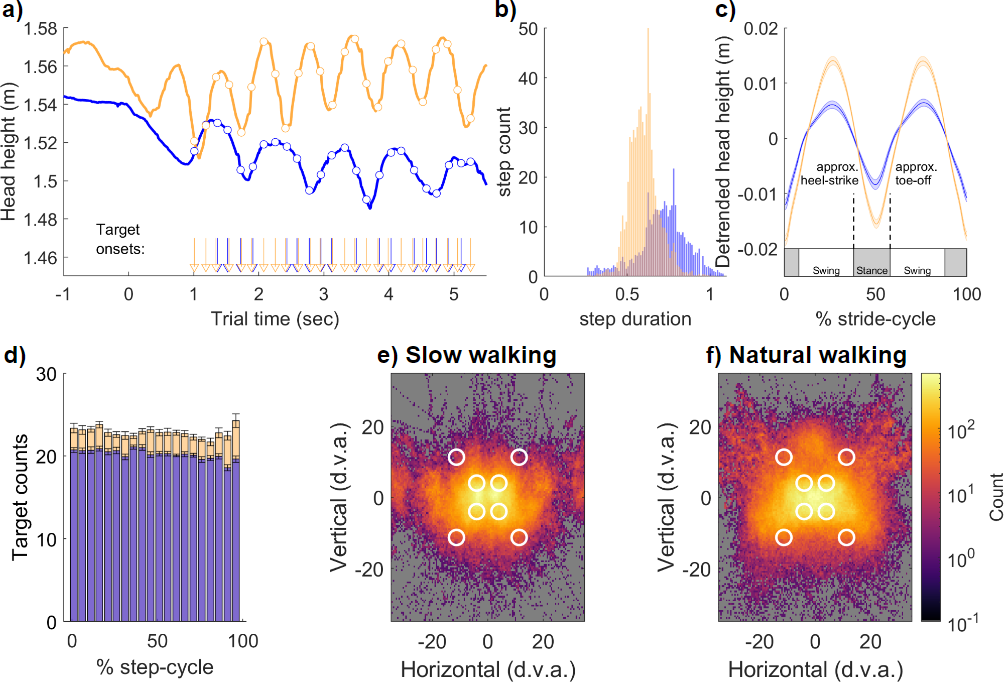
Gait extraction and gaze direction at slow and natural walking speeds. **A)** Example trials from one participant at the slow (blue) and natural (yellow) walking speeds. Raw head height is displayed, from which the peaks and troughs are used to define each step. **B)** Grand mean histograms for all step durations across participants (N=19). Slow walking conditions resulted in steps of longer duration and reduced head-height oscillation. **C)** To allow pooling of the data and comparison between walking speeds, each stride (two steps) was resampled to a normalised range of 1–100%. **D)** Target presentation density was approximately uniform over the step cycle. **E-F)** Heat maps of recorded eye directions averaged across participants for the slow and natural walking conditions. White circles note the approximate location of targets, which were at a fixed egocentric location.

### Locomotion entrains saccade onsets

We combined our gait-resampling procedure with a microsaccade detection algorithm to assess the likelihood of saccade onset with respect to the phase of the ongoing stride cycle. Clear oscillations in saccade onset timing are apparent at the group level at both slow and natural walking speeds, with saccade likelihood peaking in the approximate swing phase of each step, immediately after mid-stance. This is clear in Figures 3D-E which display the group average results (*N*=19) together with a curve displaying the best-fitting first-order Fourier model (see Methods, Equation 1).

**Figure 3.**
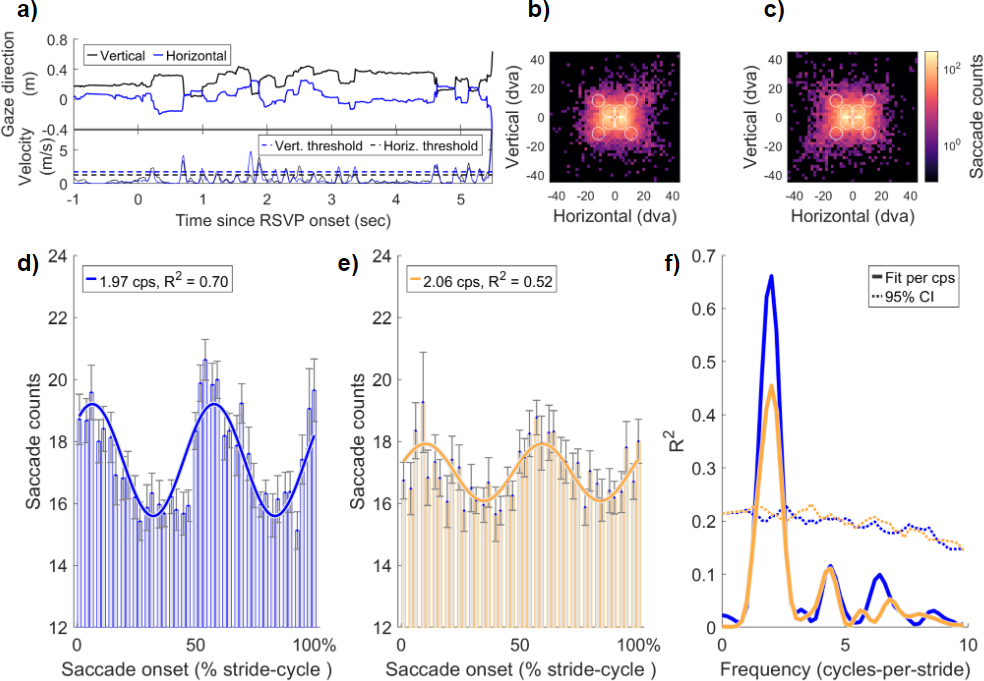
Example saccade detection, saccade distribution and saccade features time-locked to the stride cycle. **A)** Example gaze direction (black = vertical, blue = horizontal) from one trial. The bottom panel displays the same time-series converted to velocity, and the thresholds for saccade detection on the horizontal and vertical axes (see Methods). **B)** Distribution of all saccade landing positions when walking slowly and **C)** walking naturally. Approximate centre locations of the targets are shown with white circles. **D-E)** Group-level data of saccade onset timing relative to position in the stride cycle. The best-fitting first-order Fourier model is shown in each figure and approximated 2 cycles per stride (i.e., the step rate). **F)** Permutation testing of the Fourier models fitted to saccade reaction time data in d-e. The observed data were fitted with a single-component Fourier model at all frequencies between 0.2 and 10 cycles per stride (in steps of .2) and goodness-of-fit (R^2^) was calculated. The dashed line shows the upper bound of the 95% confidence interval calculated by permuting the data.

Figure 3F shows the goodness-of-fit (*R*^2^) for all Fourier frequencies in the range 0.2– 10 cycles per stride (cps; in steps of .2 cps) as well as the upper bound of the 95% confidence interval (dashed line) produced by permuting the data (see Methods). Saccade onset during slow walking speeds was best-fit by an oscillation at 1.97 cps (R^2^ = .70, range 1.40–2.40 cps above the 95th percentile). Similarly, saccade onset during natural walking speed was best-fit by an oscillation at 2.06 cps (and was significant over the range 1.40– 2.40 cps).

Together, these oscillations in saccade onset indicate that eye-movements initiated during walking entrain to the rhythm of footfall, peaking immediately after toe-off and mid-stance during the stride. We return to the timing and pattern of these oscillations in saccade likelihood in the Discussion.

### Walking speed alters the topography and timing of time-frequency power

We next analysed the changes in time-frequency power recorded with simultaneous mobile EEG at slow and natural walking speeds. Consistent with prior research, we observed pronounced fluctuations in power across a range of frequencies during each step cycle (Figure 4 A-B. Note that the x-axis shows the period of a single step, not a two-step stride cycle). These changes can be summarised as a broadband increase in power around the time of footfall, with a large decrease occurring in the swing phase at each walking speed. Interestingly, although the clusters of activity spanned a broad frequency range, the strongest effects were found at lower frequencies, predominantly in the theta (3–7 Hz) and high-alpha bands (12–15 Hz).

**Figure 4.**
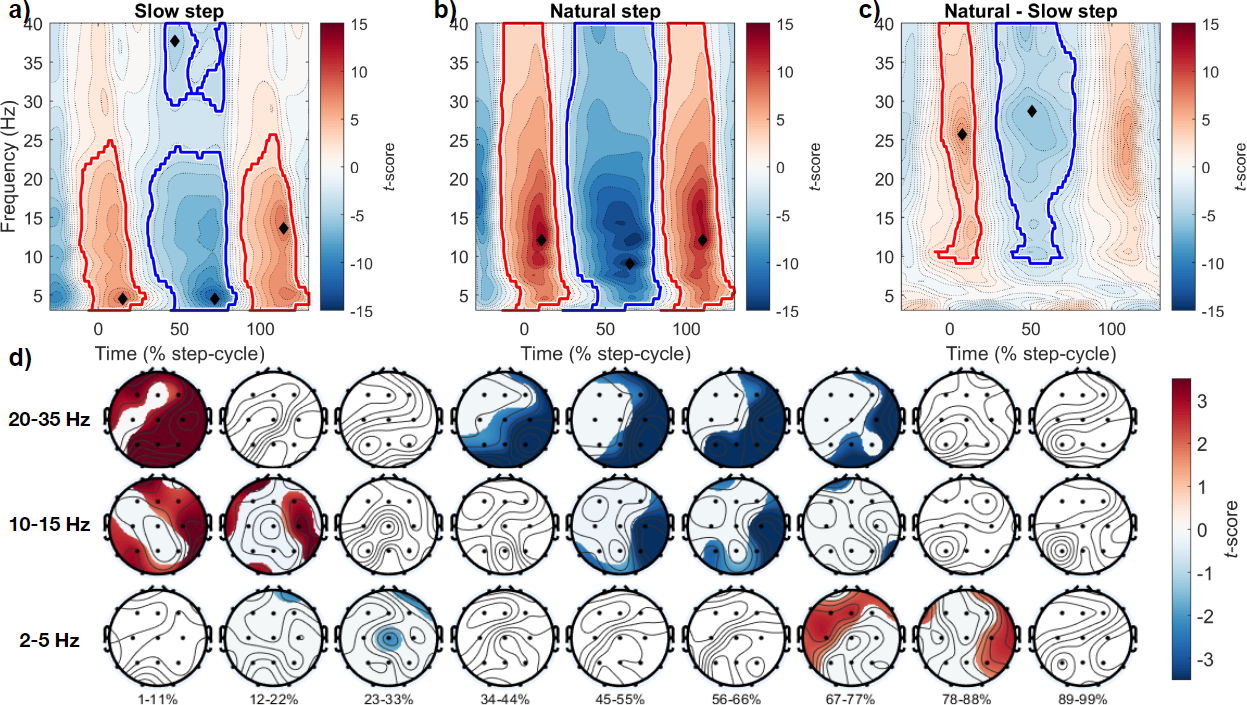
Time-frequency power over the step cycle, comparing slow and natural walking speeds. **a)** Average time-frequency power from all channels, when walking slowly. Power changes were normalised to percent relative change and significant clusters were compared to the entire epoch baseline and are displayed in each panel (see Methods). Red clusters denote increases compared to baseline, blue clusters a decrease compared to baseline. Within each cluster, the time (% step cycle) and frequency (Hz) of the peak effect are displayed with a black diamond. **b)** Time-frequency power changes during natural walking, other conventions as in a). **c)** Comparison of time-frequency activity between walking speeds (natural minus slow). **d)** Topoplots displaying the scalp distributions of differences between walking speeds averaged within specific frequency bands. The bottom, middle, and top rows show power differences at 2–5 Hz, 10–15 Hz, and 20–35 Hz, respectively. Each column represents average activity in 9 step cycle percentile bins. Significant differences (p < .05, cluster corrected) are unmasked, non-significant regions are overlaid with white masks.

More specifically, the average from all electrodes revealed that during slow walking a significant positive cluster (*p*_cluster_ <.001) increased in power, which spanned 3–24.9 Hz, during the time period ranging from −11% to 29% step-cycle completion. This difference was maximal at 4.5 Hz, 15% of the way through the step cycle (*t*(18) = 8.55). Following this, a significant negative cluster (*p*_cluster_ < .001; 3–24.1 Hz, 31–86% step cycle) occurred which was maximal at 4.5 Hz at 72% of the step cycle (*t*(18) = −11.43). A simultaneous negative cluster was present at higher frequencies during the swing phase of slow steps (*p*_cluster_ < .001; 29–40 Hz, 41–79% step cycle; max at 38 Hz, 47% step cycle; *t*(18) = −4.31). A final significant positive cluster was also present approaching the time of footfall (*p*_cluster_ < .001; 3– 26.6 Hz, 88–130%), that was maximal at 13.6 Hz (115% step cycle; *t*(18) = 8.74).

Similar results were obtained for the time-frequency dynamics during natural walking, with sequential positive (*p*_cluster_ < .001; 3–40 Hz, −15–28% step cycle, max at 12 Hz, 11% step cycle, *t*(18) = 14.18), negative (*p*_cluster_ < .001; 3–40 Hz, 23–84% step cycle, max at 9 Hz, 65% step cycle, *t*(18) = −14.68) and positive clusters (*p*_cluster_ < .001; 3–40 Hz, 84–130% step cycle, max at 12 Hz, 110% step cycle, *t*(18) = 13.61).

We also calculated the difference in time-frequency power between natural and slow walking conditions. This analysis revealed significant differences in higher frequency ranges, specifically a positive cluster at step onset (*p*_cluster_ < .001; 8.2–40 Hz, −13–21% step cycle, max at 25.6 Hz, 8% step cycle, *t*(18) = 7.03), a negative cluster during the swing phase (*p*_cluster_ < .001; 8.2–40 Hz, 24–80% step cycle, max at 28.8 Hz, 51% step cycle, *t*(18) = - 5.51), and positive cluster in the approach to heel-strike (*p*_cluster_ < .001; 9.8–40 Hz, 94–118% step cycle, max at 23.4 Hz, 109% step cycle, *t*(18) = 6.27). Figure 4A-C displays a summary of this data.

After observing strong modulations in both low and high frequency ranges in the channel-averaged data, we proceeded to investigate the spatial distribution of time-frequency power within specific frequency bands over the step cycle. For this analysis, we focused on the frequency ranges 2–5 Hz, 10–15 Hz, and 20–35 Hz, and averaged activity within these ranges over 9 step cycle percentile bins (e.g. 1–11%, 12–22%, …, 89–99% step cycle completion). Figure 4D displays a summary of this data. Most noteworthy are the distinct times and regions for significant activity within each frequency band. For example, lower frequency power (2–5 Hz) is greater during natural walking than slow walking in the approach to foot-fall (67–77%, 78–88% step cycle completion) in predominantly fronto-central and fronto-temporal electrode locations. There are no significant differences between walking speeds during the swing-phase of each step, in contrast to the higher frequency bands displayed. We return to these characteristics in the Discussion.

### Saccade-locked EEG is modulated by step cycle phase and walking speed

Finally we turned to saccade-locked time-frequency activity when separated into step cycle quartiles. For this analysis, all saccades occurring in either the slow or natural walking conditions were epoched −200 to 800 ms relative to saccade onset, and the power in each frequency band was normalised to the baseline activity within each frequency across all saccade locked epochs (see Methods). Similar to the step cycle EEG described above, we observed pronounced increases and decreases in broadband time-frequency power at both walking speeds, and relative to the phase of step cycle completion.

Unlike the step cycle EEG, where the predominant differences between walking speeds in time-frequency power were at higher frequencies **(>25 Hz, cf.** Figure 4C,D), saccade locked EEG evoked changes to frequency bands at both low and high frequencies, in a manner that depended upon the phase of locomotion.

For saccade onsets in the first quarter of the step cycle (1–24%), a significant decrease in time-frequency power was observed when walking slowly, after saccade onset, which was strongest at 5.2 Hz approx 437 ms after saccade onset (*p*_cluster_ = .015, 3.8–18 Hz, 93–500 ms, max *t*(18) = −4.52). At natural walking speeds there were significant changes to time-frequency power compared to baseline, again maximal at lower frequencies. A first positive cluster around saccade onset maximal at 6.02 Hz (*p*_cluster_ < .001, 3–40 Hz, −170–107 ms, max *t*(18) = 9.24), followed by a decrease in power at 307 ms maximal at 9 Hz (*p*_cluster_ < .001, 3–40 Hz, 106–406 ms, max *t*(18) = −13.36) and a later increase in power at 6.7 Hz (*p*_cluster_ < .001, 3–34.7 Hz, 443–700 ms, max at 6.7 Hz, 620 ms, *t*(18) = 7.42). There were no differences in time-frequency power when comparing walking speeds (Figure 5A-D).

**Figure 5.**
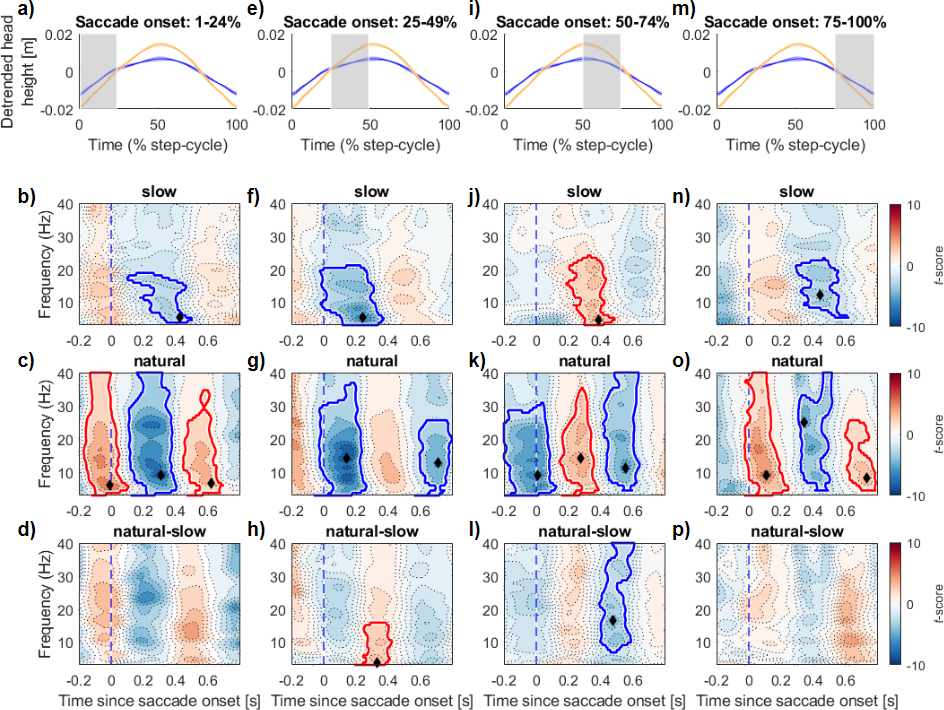
Saccade-locked EEG time-frequency dynamics during locomotion are modulated by gait-cycle phase. **A)** The average detrended change in head height over all participants (blue = slow walk speed, yellow = natural walk speed). The shaded patch denotes the region (in step cycle %) of pooled saccade onsets used in the saccade-locked EEG in b-d). **B)** Saccade-locked EEG from the first quarter of the step cycle (1–24%), significant negative cluster shown in blue, with peak frequency and time point of the effect displayed with a black diamond. **C)** Saccade-locked EEG in the same quartile, increases in time-frequency power are shown in red. **D)** The difference in time-frequency power when comparing natural and slow steps for this step cycle quartile. **E-P)** Shows the same data format as in a-d repeated over subsequent quartiles of the step cycle. All clusters corrected for multiple comparisons using non-parametric permutation tests (Maris & Oostenveld, 2007).

For saccades occurring in the second quarter of the step cycle (25–49%), a similar pattern emerged. Time frequency EEG power during slow speeds significantly decreased after saccade onset with a peak effect in the high theta range (*p*_cluster_ = .005, 3–21 Hz, - 230ms–370 ms, max at 5.2 Hz, 243 ms, *t*(18) = −8.99). At natural speeds, a first negative cluster peaked 140 ms after saccade onset (*p*_cluster_ < .001, 3–37 Hz, −6ms–270 ms, max at 14.3 Hz, 143 ms, *t*(18) = −12.45), and another at approximately 500 ms (*p*_cluster_ < .001, 3– 25.6 Hz, 560 ms–793 ms, max at 12.8 Hz, 710 ms, *t*(18) = −7.25). When comparing walking speeds, there was an increase in low frequency power during natural walking (*p*_cluster_ = .035, 3–15.8 Hz, 20ms–426 ms, max at 3.75 Hz, 330 ms, *t*(18) = 5.63) (Figure 5E-H).

When saccades occurred after the mid-point of each step cycle (50–74%) there was an increase in low-frequency power approx 383 ms after onset when walking slowly, which peaked at 4.5 Hz (*p*_cluster_ < .001, 3–24.25 Hz, 186–487ms, max *t*(18) = 6.66). At natural speeds, a decrease in power (*p*_cluster_ < .001, 3–31 Hz, −200–123 ms, max at 0 ms, 9 Hz, *t*(18) = −9.57), then increase (*p*_cluster_ < .025, 3–35.5 Hz, 143–386 ms, max at 276 ms, 14 Hz, *t*(18) = 6.88), then decrease was again present (*p*_cluster_ < .025, 4.5–40 Hz, 413–640 ms, max at 553 ms, 11.3 Hz, *t*(18) = −7.34). When comparing walking speeds, there was a decrease in time-frequency power in the natural condition which peaked 477 ms after onset at 16.6 Hz (*p*_cluster_ < .001, 6.7–40 Hz, 386–606 ms, max, *t*(18) = −5.72).

For saccade onsets late in the step cycle (75-100%), there was a significant negative cluster when walking slowly (*p*_cluster_ < .001, 5.2–23.4 Hz, 290–623 ms, max at 443 ms, 12 Hz, *t*(18) = −5.21). At natural speeds, there was again a sequence of positive (*p*_cluster_ < .001, 3–40 Hz, 3–250 ms, max at 106 ms, 9 Hz, *t*(18) = 9.27), negative (*p*_cluster_ < .001, 4.5–40 Hz, 305–517 ms, max at 343 ms, 24.9 Hz, *t*(18) = −8.54), then positive clusters (*p*_cluster_ < .001, 4.5–25.6 Hz, 497–793 ms, max at 730 ms, 8.3 Hz, *t*(18) = 5.81).

The overall pattern of these results can be summarised as an increase in modulation of time-frequency power when walking at natural speeds. A clear pattern of sequential positive and negative clusters is also evident – with the order of these clusters driven by the relative phase of the step cycle. Also noteworthy are the frequencies at which these effects are maximal – predominantly in the low frequency band when walking slowly (average frequency of peak effects 6.73 Hz; *SD* = 3.5 Hz), and slightly higher at natural walking speeds (M= 11.34 Hz, *SD* = 5.29).

## Discussion

In this study we measured eye movements and EEG activity in observers doing a visual oddball detection task. A critical additional feature was that our observers were active while the measurements were being made, walking back and forth along a path at two walking speeds. Doing the task while engaged in natural active behaviour is important because we are fundamentally active creatures, constantly reaching, head-turning, walking and moving our eyes as we engage in our daily routines and complete relevant tasks. It is therefore important to study how active behaviour might alter cognitive and perceptual performance, and how one type of action might influence another, for a full understanding to be achieved. With this principle in mind, we designed the current experiment around a walking participant using new advances such as mobile EEG and wireless VR with integrated eye movement recordings and three-dimensional position tracking.

Our results confirm that important differences do emerge in behaviour and neural activity when measured during action. Specifically, we showed that both the likelihood of making saccades and EEG time-frequency power are modulated rhythmically over the gait cycle, producing an oscillation that matched the step rate (approximately 2 Hz). Moreover, the saccadic and EEG oscillations show a consistent phase relationship with the gait cycle (Figures 3D,E & 4A,B), indicating both are linked to gait. By comparing saccade onsets and EEG at two walking speeds, we confirmed that saccadic and EEG oscillations are linked to the phase of the stride-cycle by showing they entrain to the rhythm of footfall even at a slower pace. We also observe separable dynamics to the time-frequency power simultaneously recorded during walking at different speeds, which provide strong considerations for interpreting saccade-locked EEG during locomotion, and mobile-EEG in general.

### Saccade onsets entrain to the rhythm of footfall

Participants in our task were required to walk in a straight line while fixating a central cross, and monitored eight peripheral target locations during an oddball task. Targets appeared every 200 ms (∼ 5 Hz), with onsets that were approximately evenly distributed across the step cycle (Figure 2D). Despite this uniform presentation rate we observed strong modulations to the likelihood of saccade onsets that closely matched the step cycle frequency, with a period of approximately 2 cycles per stride.

The timing of these phasic changes to the likelihood of saccade onset are noteworthy for several reasons. Despite the subjectively smooth and continuous experience of walking, the rhythm of walking is supported by ballistic periods of heightened sensory demand. For example, in the swing phase immediately after toe-off, the predictability of head-movements are at their lowest (MacNeilage & Glasauer, 2017), leading to increased vestibular demand in order to maintain balance and posture (Bent et al., 2004; Dakin et al., 2013). Similarly, recent work tracking eye-movements during movement through challenging terrain has shown an increase in gaze density for locations approximately 2-steps ahead (Bonnen et al., 2021; Matthis et al., 2018; Matthis & Fajen, 2014), and that information about the terrain must be viewed within a critical window (prior to heel-strike) for smooth locomotion to continue (Matthis et al., 2017; Matthis & Fajen, 2013). These changes in demand coincide with the period of heightened saccade onset probability that we report. Although in our task saccades were not necessary only for smooth locomotion, it is likely that their occurrence helps to orient the body in an uncertain environment to guide locomotion. Indeed, we observed an increased modulation in the strength of saccade onset probability when walking slowly – a task which is known to place additional demands on sensory and cognitive systems (Akaiwa et al., 2022; Al-Yahya et al., 2011; Lajoie et al., 2016). Future research may further increase sensory uncertainty to investigate these modulations in saccade onset probability.

### Distinct time-frequency power changes that depend on walking speed

We observed significant changes in time-frequency power that were also coupled to the phases of locomotion. Previous examinations of mobile-EEG activity have noted similar large-scale fluctuations in power. In the first study to report intra-stride fluctuations of electrocortical activity during walking, Gwin et al (2011) identified broadband and sequential increases and decreases in event-related spectral perturbations. Using a high-density EEG array and guided ICA, these fluctuations were spatially localised to anterior cingulate, parietal and sensorimotor cortex, with no differences between activity at two speeds of treadmill walking (0.8 and 1.25 m/s). We also observed strong fluctuations in alpha and beta-band activity that were phase-locked to the procession of the step cycle, which unlike this previous research significantly differed at each walking speed. More specifically, the fluctuations in power were maximal in different frequency bands depending on walking speed, peaking in the approximate theta band when walking slowly, and approximate alpha band at natural walking speeds (Figure 4).

The frequency of these peak effects are noteworthy for several reasons. As has been noted in the past (Castermans et al., 2014; Gramann et al., 2011, 2014; Gwin et al., 2010), it is critical to remove motion signals and potential artefacts prior to interpreting underlying scalp activity, particularly with respect to activity in the low delta and high-gamma range, for contamination due to the step-rate and muscle-activity respectively (Castermans et al., 2014). Our procedure incorporated highly conservative exclusion criteria and stringent artefact removal, and observed strongest effects outside the frequency of the step-rate (> 2 Hz), and below high-gamma fluctuations scrutinised in prior research (<50-150 Hz). When analysing the topography of this frequency-band activity, we also observed a relative increase in frontal theta-band activity at natural walking speeds, just prior to heel-strike (Figure 5D). This change in frontal theta is reminiscent of prior reports examining reactive changes to balance and posture, where it has been proposed that theta dynamics may represent a process of action monitoring to maintain balance (Stokkermans et al., 2022).

### Saccades and EEG

Lastly, we viewed changes in the EEG signal around saccades, when those saccades were early or late in the step cycle, during natural and slow walking. We observed dynamic changes in the low to mid frequency bands, depending on whether saccades co-occurred with each of four parts of the step, from onset to footfall. We found that saccade-locked EEG power increased during swing, and decreased during approach to heel-strike, more strongly when walking at natural speeds compared to slow. Critically, these walking speed effects centred on low to mid frequencies, peaking at 3.75 Hz for swing and 16.6 Hz for approach to heel-strike. This contrasts with what we observed in the EEG when accounting for step cycle alone, where walking speed effects were maximal between 20 and 30 Hz.

Given the practical challenges of recording eye movements and EEG during free movement, it is unsurprising that there are few comparable studies. However, static work relating eye movements to EEG highlights their close relationship. Liu et al. (2023) show changes in ipsilateral alpha power during microsaccades, arguing that even very subtle shifts in gaze can alter signals associated with visual and spatial cognition. This relationship also appears to go in the other direction. Staudigl et al. (2017) used magnetoencephalography (MEG), a close cousin to EEG, to show that posterior alpha phase predicts saccades during memory encoding. That is, saccades may both prime changes in low frequency power, and be triggered by them, in regions that are critical for interpreting our visual world. Here, we show that saccade-related changes in EEG interact with walking speed, and that we can observe this over and above step-cycle-related changes. Together, the co-registration of eye and body movements with EEG demonstrates a way towards understanding motor and perceptual processes in rich neural data during natural movement.

## Conclusion

Overall, our results make an important general point in showing that extrapolating knowledge obtained from traditional experiments to predict outcomes in richer, active environments is not a valid extension. The current knowledge base was very largely built on data from passive observers sitting immobile in darkened laboratories who respond to simple stimuli without the distraction of real-world clutter and the competing demands of being active. This approach has undoubtedly been a rich and successful one but has led to an artificially narrow understanding. On this view, we would not expect that eye movement and EEG activity, as well as visual performance on the oddball task, would show oscillations at the step rate when measured in a walking participant. It is indeed rather surprising to find oscillatory results given that the visual task was an uninterrupted one that required continual monitoring and that walking seems a near-effortless constant activity. Yet, despite this, clear modulations were found, all linked to the gait cycle.

These results have important implications. 1) Because locomotion imposes a rhythmic oscillation on neural, saccadic and perceptual activity, it therefore creates good and bad phases for perceptual performance and saccadic behaviour. Saccades, for example, are more likely to occur at certain moments than others, peaking just after footfall. It likely this temporal clustering of eye movements would have consequences for tasks such as visual search and change detection and this could be explored in future experiments. Similarly, the rhythm of walking imposes an oscillation on brain activity which needs to be taken into account when analysing EEG data. 2) A second, more general, implication is to underscore the importance of conducting experiments in active observers in more realistic contexts. The technical hurdles that previously made such experiments very challenging to implement have largely been overcome and we can now co-register multiple information streams such as EEG and eye movements and various biomarkers in active observers behaving in rich virtual or real environments. Experiments conducted along these lines will add new layers to our knowledge base as more active and realistic features are added to experiments which will, in turn, help bridge the gap between lab-based experiments and real-world applications.

